# Single-cell ATAC-seq Reveals OVOL2 as a Downstream Negative Regulator of PRL-Mediated Chromatin Accessibility

**DOI:** 10.64898/2026.04.01.715828

**Authors:** Nelmari Ruiz-Otero, Jin-Yong Chung, Ronadip R. Banerjee

## Abstract

Maternal pancreatic β-cells undergo functional and structural changes to adapt to increased metabolic demands during pregnancy. Lactogen signaling via the prolactin receptor (PRLR) contributes to these adaptations by increasing β-cell mass, insulin transcription and glucose-stimulated insulin secretion[1–4]. In other lactogen-responsive tissues such as the mammary glands and specific hypothalamic nuclei, gestation induces epigenetic changes, some of which persist long after birth[5, 6]. We have previously found that prolactin treatment in islets regulates the expression of epigenetic modifiers[7, 8]. However, whether lactogen signaling in β-cells mediates epigenetic changes to regulate chromatin accessibility has not been examined. Therefore, our objective was to determine whether PRLR signaling alters chromatin accessibility of β-cells to facilitate transcriptional regulation.

Using single-cell ATAC-sequencing, we identified differentially accessible regions (DARs) in β-cells which had 718 overrepresented motifs following prolactin treatment of murine islets. Validating this approach, these included motifs bound by established PRLR signaling effectors such as the STAT family of transcription factors (TFs). Using RNA-sequencing we identified transcriptional changes in 41 TFs whose motifs were overrepresented in DARs, including several previously linked to PRLR signaling within β-cells, including *Myc*, *Mafb* and *Esr1*. Importantly, we also identified TFs not previously associated with PRLR signaling, including OVOL2 an established regulator of epigenetic landscape within cells. OVOL2 is a transcription factor involved in EMT inhibition and energy homeostasis with unknown roles in pancreatic β-cells. Here, we establish that OVOL2 acts as a negative regulator of lactogen-dependent effects on β-cell proliferation, establishing a novel regulator of PRLR signaling.

## Introduction

Hormone physiology in pregnancy is both intricate and dynamic. During pregnancy the female body undergoes multiple adaptations to increase insulin secretion[9]. Failure to adapt results in gestational diabetes (GDM), conferring increased short– and long-term health risks to the offspring, and profoundly affecting maternal health[10, 11]. Although we have a broad understanding of how GDM affects the progeny’s health long-term, the pathophysiological effects of GDM in maternal health long after birth are well less understood, displaying a knowledge void in a disease affecting around one in seven pregnancies worldwide[12, 13].

Prolactin receptor (PRLR) signaling is a critical pathway regulating the insulin-producing β-cells during pregnancy. Placental lactogens and the pituitary gland-derived prolactin (PRL) signal via PRLR expressed by the β-cells where it acts via JAK2/STAT5[14–16] and other downstream signaling effector pathways to expand β-cell mass, increase insulin expression and enhance glucose-stimulated insulin secretion (GSIS)[3, 17, 18]. We have previously established that pregnant female mice with a conditional deletion of PRLR in β-cells (β-PrlrKO mice) exhibit GDM, displaying normoglycemia before pregnancy but develop hyperglycemia in late gestation along with a lack of β-cell mass expansion, illustrating a critical requirement for PRLR signaling in gestational adaptation[19]. Therefore, it is crucial to understand the underlying mechanisms by which PRLR signaling acts on β-cells, as defects in these processes can lead to GDM.

Other lactogen-responsive tissues, such as the mammary glands and some hypothalamic nuclei involved in energy balance, display epigenetic responses to pregnancy, many which persist after birth[5, 6]. In the mammary glands, PRLR signaling induces milk production and secretion, similar to how it induces insulin biosynthesis and secretion in β-cells [20]. Additionally, mammary involution, which occurs when circulating lactogens diminish, mirrors how β-cell mass recedes and insulin levels return to a “non-pregnant state” when lactation is terminated[20]. Due to these similarities, we hypothesized that lactogen activation of PRLR signaling might similarly induce epigenetic modifications changing chromatin accessibility within β-cells.

To deconvolute the direct role of PRLR signaling amongst the complex hormonal landscape during gestation we used a combination of *ex vivo* primary islet cell assays and a murine cell culture model of β-cells. Here, we show that PRLR signaling *ex vivo* modulates chromatin accessibility, revealing specific motif signatures of previously uncharacterized PRLR-dependent TFs activation within β-cells. Among them, OVOL2 plays a significant role in regulating PRLR-dependent transcription of genes involved in β-cell homeostasis. OVOL2 is a transcription factor from the OVO family, whose homolog *ovo* was first characterized in Drosophila as a sex determination gene[21]. OVOL2 is primarily associated with repression of the epithelial to mesenchymal transition (EMT), mainly by TGFβ signaling inhibition[21–23]. OVOL2 has also been identified as a novel regulator of energy homeostasis. For example, it plays an inhibitory role in glucose import, regulates aerobic respiration, limits adipogenesis, and hypomorphic mutations of the *Ovol2* gene result in impaired glucose tolerance and reduced insulin sensitivity[24–26]. Furthermore, OVOL2 is known to regulate chromatin accessibility by recruiting NCoR and histone-modifying enzymes (HDACs)[21–23]. Previous studies have identified OVOL2 as a direct target of PDX1, a TF critical in regulating β-cell identity and function[27]. However, OVOL2’s importance in β-cells, outside of EMT gene regulation, had not been elucidated.

Our studies show that OVOL2 is expressed by β-cells during gestation in a PRLR-dependent manner. Overexpression of OVOL2 impairs PRL-mediated proliferation, suggesting a previously unrecognized role for OVOL2 as a PRLR signaling regulator, and for PRLR signaling to regulate β-cell functions and adaptations through structural modification of chromatin.

## Results

### PRLR signaling in islet cells induces changes in chromatin accessibility

PRLR signaling mediates many intracellular processes in β-cells to contribute to gestational adaptations. Due to the complex hormonal and metabolic landscape during gestation, we turned to a reductionist model, in which *ex vivo* treatment of islets using recombinant mouse PRL induces transcriptional responses parallel to those observed in late mouse gestation[28]. To determine whether activating PRLR signaling affects chromatin accessibility, we performed scATAC-sequencing in the presence or absence of PRL on islets isolated from virgin female mice 8-12 weeks of age from either wild type C57BL/6 (WT) or a mice lacking PRLR specifically in β-cells, β-PrlrKO (*RIP-Cre; Prlr^f/f^*)[19] **(Figure 1a, Supplemental fig. 1a, and GSE319576)**. PRL treatment of cultured islets in vitro for 24 hours exhibited transcriptional responses like those observed *in vivo* [1, 18, 29–34], including the upregulation of *Tph1* and *Tph2* **(Supplemental fig. 1b)** which code for the enzymes responsible for serotonin synthesis.

**Figure 1:**
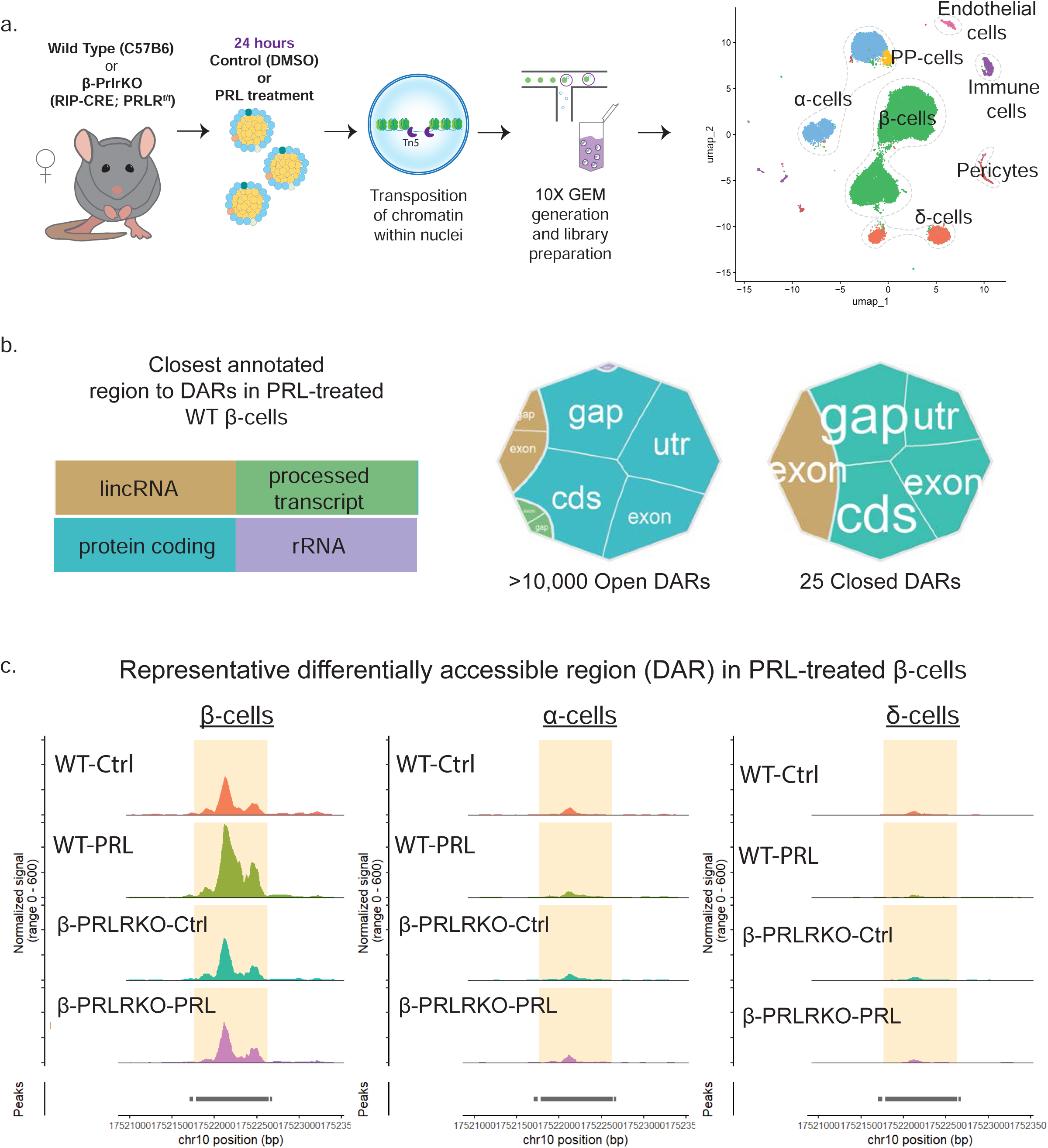
scATAC-sequencing of islets show that PRLR signaling mediates DARs in β-cells. a. Experimental outline of scATAC-sequencing from islet cells treated with PRL b. Tree map of closest genomic features annotated to the differentially accessible regions observed among WT β-cells after PRL treatment. c. Representative coverage plot PRL-dependent DARs in WT β-cells but not in β-PrlrKO β-cells or WT α– and δ-cells

To assess the specific effect of PRL treatment in β-cells **(Supplemental fig. 1c)**, we compared chromatin accessibility from WT non-treated versus PRL-treated cells. We found over 10,000 differentially accessible regions (PRL-DARs), the majority of which showed increased accessibility (Open DARs). When we assessed the closest annotated feature of PRL-DARs, the majority fell in the vicinity of protein coding regions **(Figure 1b and Supplemental fig. 1d)**. Consistent with the CRE-dependent recombination, β-PrlrKO β-cells, lacked chromatin accessible sites in exon 5 of the *Prlr* gene **(Supplemental fig. 2a)**. In contrast to WTs, PRL-treated β-cells from β-PrlrKO had 2832 PRL-DARs all showed decreased accessibility. Interestingly, vehicle treated WT and β-PrlrKO β-cells showed some basal chromatin accessibility differences both in open and closed regions **(Supplemental fig. 2b)**. We previously reported no glucose metabolic alterations in virgin β-PrlrKO females, but these basal chromatin accessibility differences suggest that further examination may reveal subtle functional or transcriptomic differences in non-gestational β-cells lacking PRLR under specific conditions. In contrast to WT β-cells, WT α– and δ-cells displayed only 537 and 12 PRL-DARs respectively, consistent with their lack of PRLR expression **(Figure 1c)**. Thus, consistent with prior studies, β-cells are the main targets of lactogen signaling within murine islets[3, 19, 35]. ***Together, we present the first direct evidence of PRLR-mediated chromatin remodeling within islet cells*.**

To examine the relationship of chromatin changes with transcriptional regulation, we performed RNA-sequencing on a subset of the WT islets treated simultaneously to those in the scATAC-seq experiment described above. We identified over 1000 differentially expressed genes (PRL-DEGs), of which only 303 were downregulated **(GSE319525).** Consistent with previously reported data, PRL-treated islets had increased transcripts of *Tph1*, *Tph2, Cish*, and *Prlr,* **(Supplemental fig. 3a)** which are regulated in a PRLR-dependent manner during gestation[19]. Importantly, we found that a subset of genes associated with PRLR functions within islets, such as *Cish,* which is a negative regulator of PRLR [36], and *Enpp2* which promotes β-cell proliferation[37], contained PRL-DARs **(Figure 2a)**, defining a critical link between chromatin structure and gene expression driven by PRLR signaling. In contrast, other PRL-DEGs, such as *Tph1* and *Tph2* had no changes in chromatin accessibility of the gene and proximal promoter, nor their immediate chromosomal vicinity **(Figure 2b)**, although we cannot formally exclude PRLR-mediated chromatin changes to *Tph* expression, as trans and long-distanced enhancers driven by PRLR might modulate their expression[38].

**Figure 2:**
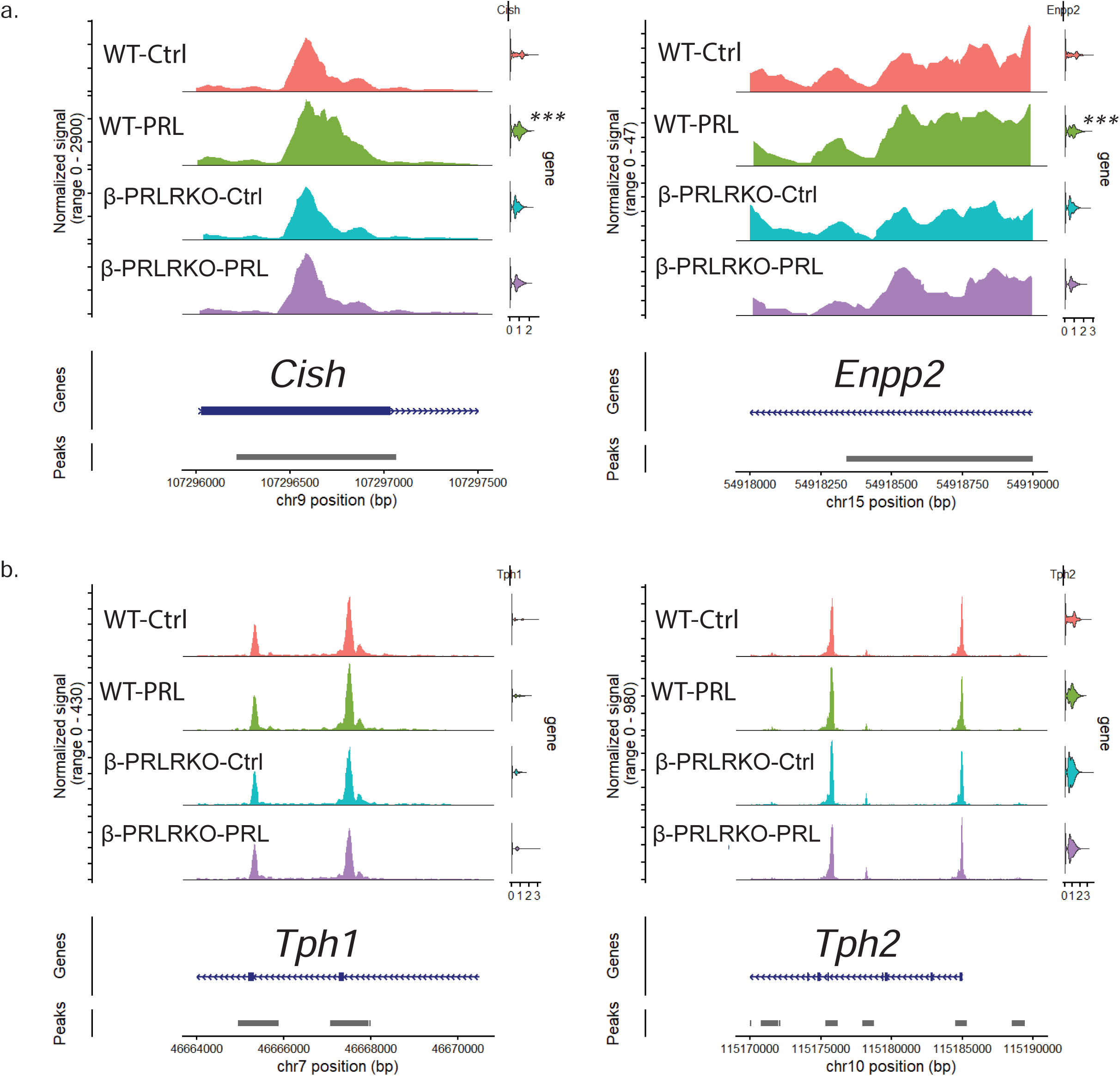
Visualization of chromatin accessibility within PRL-DEGs. a. Coverage plots from β-cells showing *Cish* and *Enpp2* differential chromatin accessibility in WT β-cells treated with PRL b. Coverage plots from β-cells showing *Tph1* and *Tph2* similar chromatin accessibility in β-cells regardless of genotype or treatment

### Identification of novel TFs involved in PRLR signaling in β-cells

We next sought to determine whether PRL-DARs had enrichment of TF motifs from which we could identify TFs regulating chromatin changes downstream of PRLR signaling. Chromatin accessibility is regulated by histone modifications and chromatin methylation directed by TFs[38]. We first looked for overrepresented motifs present in PRL-DARs within WT but not in β-PrlrKO β-cells. This list of motifs included those bound by the STAT family of TFs **(Supplemental fig. 3b)**, which are well established PRLR-signal transducers[33]. Of the 718 motifs identified, we decided to pursue overrepresented motifs of TFs that were also differentially expressed in response to PRL **(Figure 3a).** Validating this approach, these PRL-enriched TFs include some TFs we and others have previously identified as PRLR mediators, such as *Myc*, *Esr1, Jun* and *Mafb* [7, 19, 29]. Among these, OVOL2 had the highest motif enrichment score and fold change in transcription and had not been previously linked to β-cell functions in any context. In addition to OVOL2, we identified other TFs not previously implicated in PRLR signaling within β-cells, including GLI3, ZBTB7C and ALX4, TFs whose role outside the islets range from regulating developmental genes to fatty acid metabolism and cellular structure[39–41]. We focused on OVOL2 as a PRL-dependent chromatin regulator because OVOL2 has been previously reported as a chromatin modifier regulator[21, 23]. Confirming our RNA-seq finding, PRL-dependent increased OVOL2 expression in WT islets following PRL stimulation by qPCR analyses **(Figure 3b). *In sum, we have identified novel TFs not previously associated with PRLR signaling or β-cell functions, including Ovol2, which is induced by PRL in cultured islets*.**

**Figure 3:**
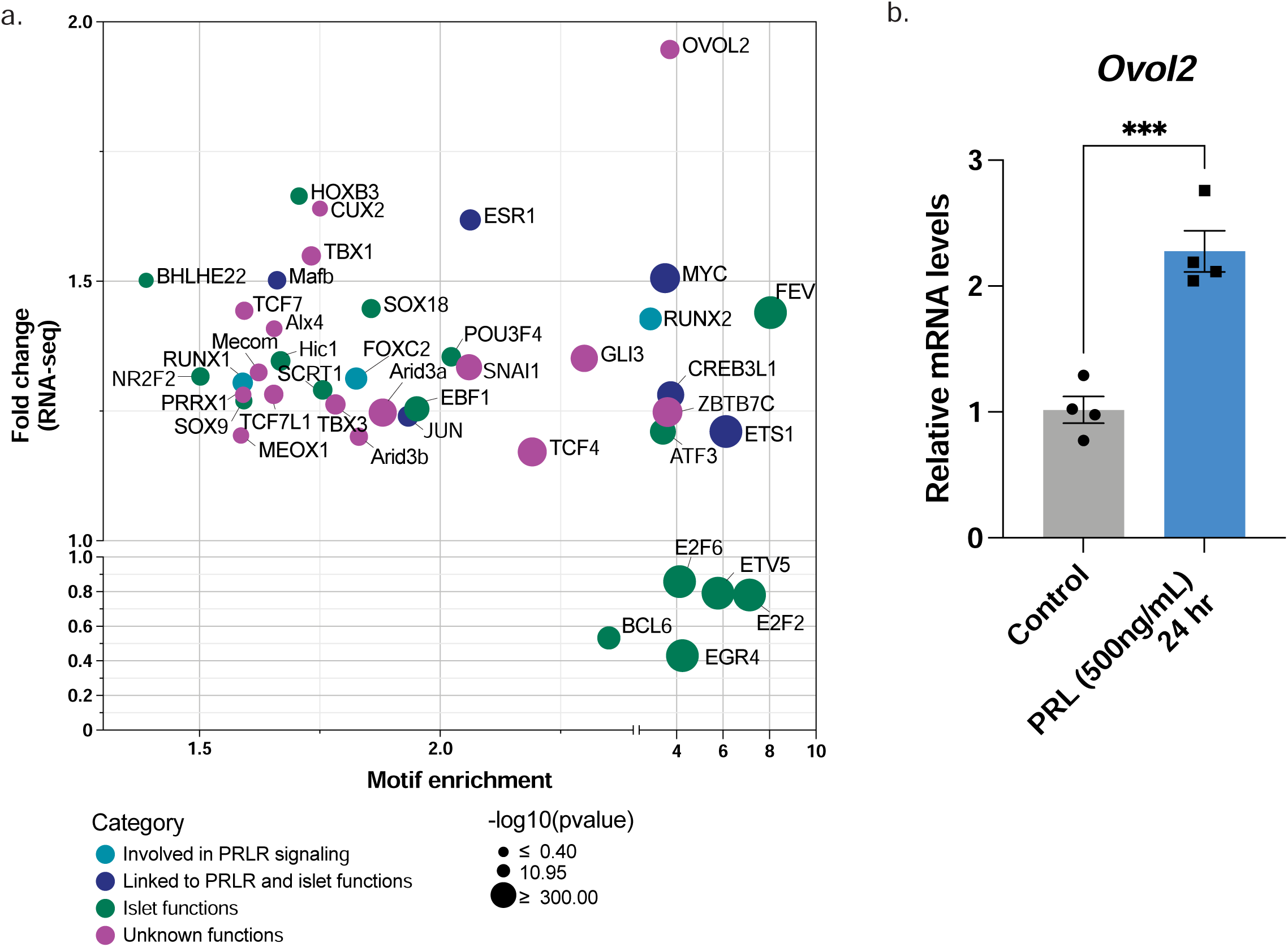
Identification TFs which are PRL-DEGs and with overrepresented motifs in WT β-cells PRL-DARs. a. Bubble plot to visualize TFs regulated transcriptionally by PRL treatment and whose motifs are overrepresented in β-cell PRL-DARs. Motif enrichment score from β-cells calculated from PRL-DARs and shown in the x-axis. Fold change calculated from RNA-seq is shown in the y-axis. Size of the bubbles is dependent on the p value calculated for motif enrichment and colors are assigned based on category (See Supp fig 4b). b. qPCR of *Ovol2* after PRL-treatment. Relative mRNA levels were normalized to *Rps29* and calculated using the 2^-ΔΔCT^ method. Student t-test: *** p value <0.001.

### PRLR-dependent regulation of OVOL2 binding and functions

OVOL2 has been implicated as a canonical TF enhancing and repressing gene expression and regulating chromatin access through interactions with histone-modifying enzymes (HDACs) and the NCoR complex [21, 23]. Therefore, we hypothesized that PRLR signaling regulated OVOL2 target binding and functions within islet cells thereby impacting chromatin structure. We compared a previously reported list of OVOL2 targets identified in other tissues, such as mammary epithelium, keratinocytes, and thyroid carcinoma cell models[21], with our RNA-seq results of PRL-treated islets to identify potential PRLR-dependent OVOL2 gene targets in β-cells. We designed primers for the promoter region of the selected genes, including regions containing OVOL2 predicted binding sites according to the JASPAR database[42]. Using chromatin immunoprecipitation followed by quantitative PCR (ChIP-qPCR) in islets treated with PRL, we confirmed PRL-dependent binding of OVOL2 to the promoter regions of *Snai1*, a well-established Ovol2 target in other tissues and which functions to *s*uppress EMT, *Rhoj,* which is involved in restraining mitosis, *Myc* which is a TF with established roles in β-cell proliferation and survival **(Figure 4a),** and *Jun,* **(Supplemental fig. 4a),** which we have shown functions to promote β-cell survival in certain contexts[29].

**Figure 4:**
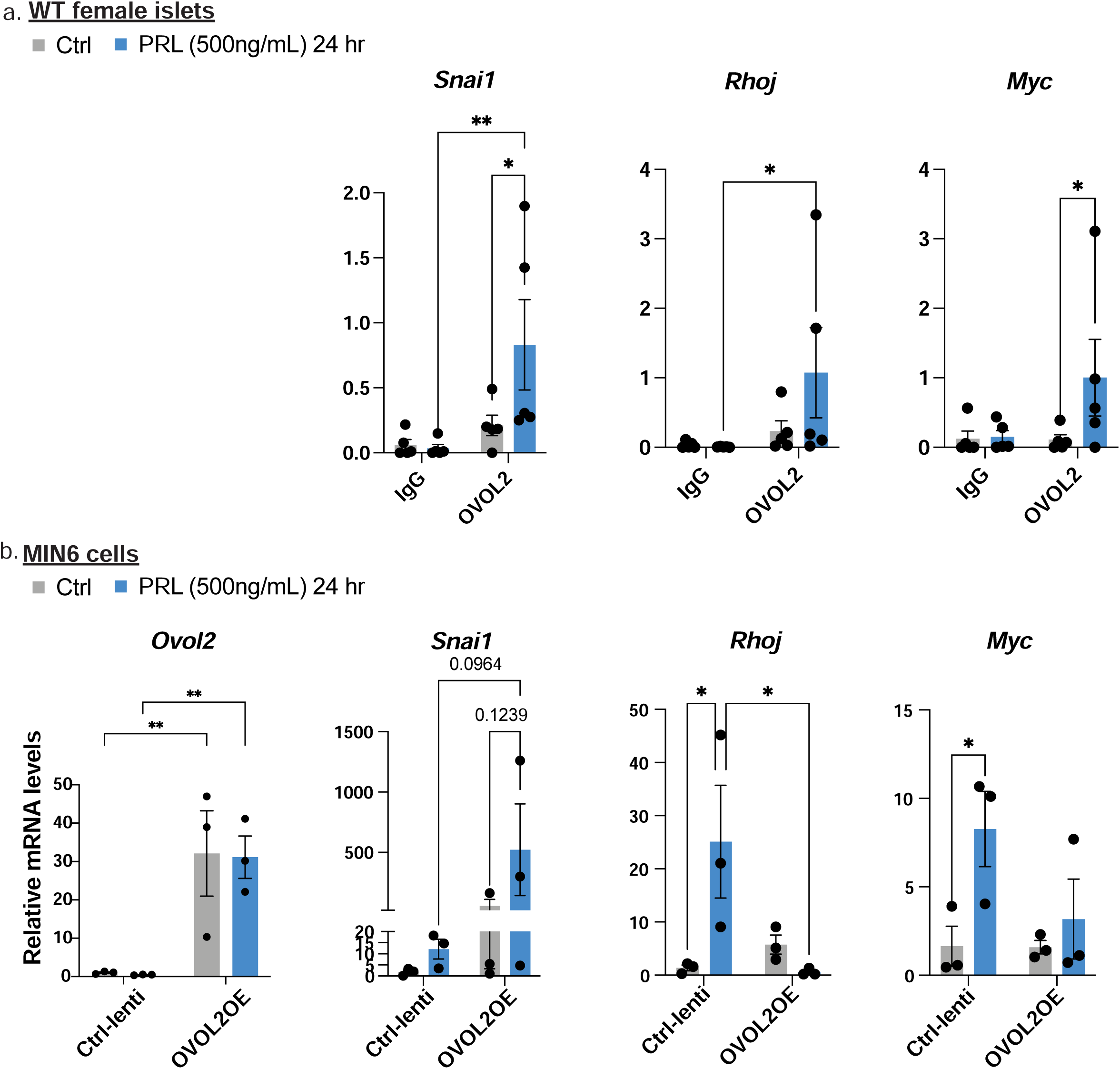
PRL-dependent binding and gene regulation by OVOL2. a. ChIP-qPCR from islets treated with PRL. Binding to promoters was calculated in relation to input (% input) and evaluated against rabbit IgG as a control for non-specific binding. Means ± SEM for N= 5 experiments: 2Way ANOVA: * p-values <0.05 and ** p-values <0.01. b. qPCR evaluation of MIN6 cells infected with control (Ctrl-lenti) or OVOL2OE after PRL treatment. Relative mRNA levels were normalized to *Rps29* and calculated using the 2^-ΔΔCT^ method. Means ± SEM for N= 3 plates per condition and treatment combination. 2Way ANOVA: p-values <0.2 shown, * p-values <0.05 and ** p-values <0.01.

To further discern the specific functions of OVOL2 we turned to a murine cell culture model of β-cells, MIN6 cells[43], which retain PRL-responsiveness as evidenced by increased *Tph1* and *Tph2* expression **(Supplemental figure 4b).** We performed lentiviral infection of MIN6 cells with a custom vector to overexpress GFP-tagged OVOL2 (VectorBuilder: VB250505-1105ZYC, hereafter OVOL2OE). OVOL2OE MIN6 cells showed a ∼30-fold increase in *Ovol2* transcription **(Figure 4b)**, regardless of PRL treatment. There was a noticeable trend toward increased *Snai1* transcription in OVOL2OE cells which was enhanced by PRL treatment that did not achieve statistical significance **(Figure 4c).** However, contrary to our initial model, PRL-dependent induction of *Myc* and *Rhoj* was prevented by OVOL2 overexpression. These results led us to hypothesize that OVOL2 might act as a negative regulator of PRLR signaling constraining signaling amplification after lactogen withdrawal.

To assess whether OVOL2 impaired PRLR signaling effects, we assessed MIN6 cell proliferation by BrdU incorporation in the presence of PRL. Control MIN6 cells showed increased proliferation in response to PRL treatment as expected. However, PRL treatment failed to enhance proliferation of OVOL2OE cells (**Figure 5a and 5b**), indicating that OVOL2 overexpression negatively regulates PRLR signaling-mediated proliferative effects.

**Figure 5:**
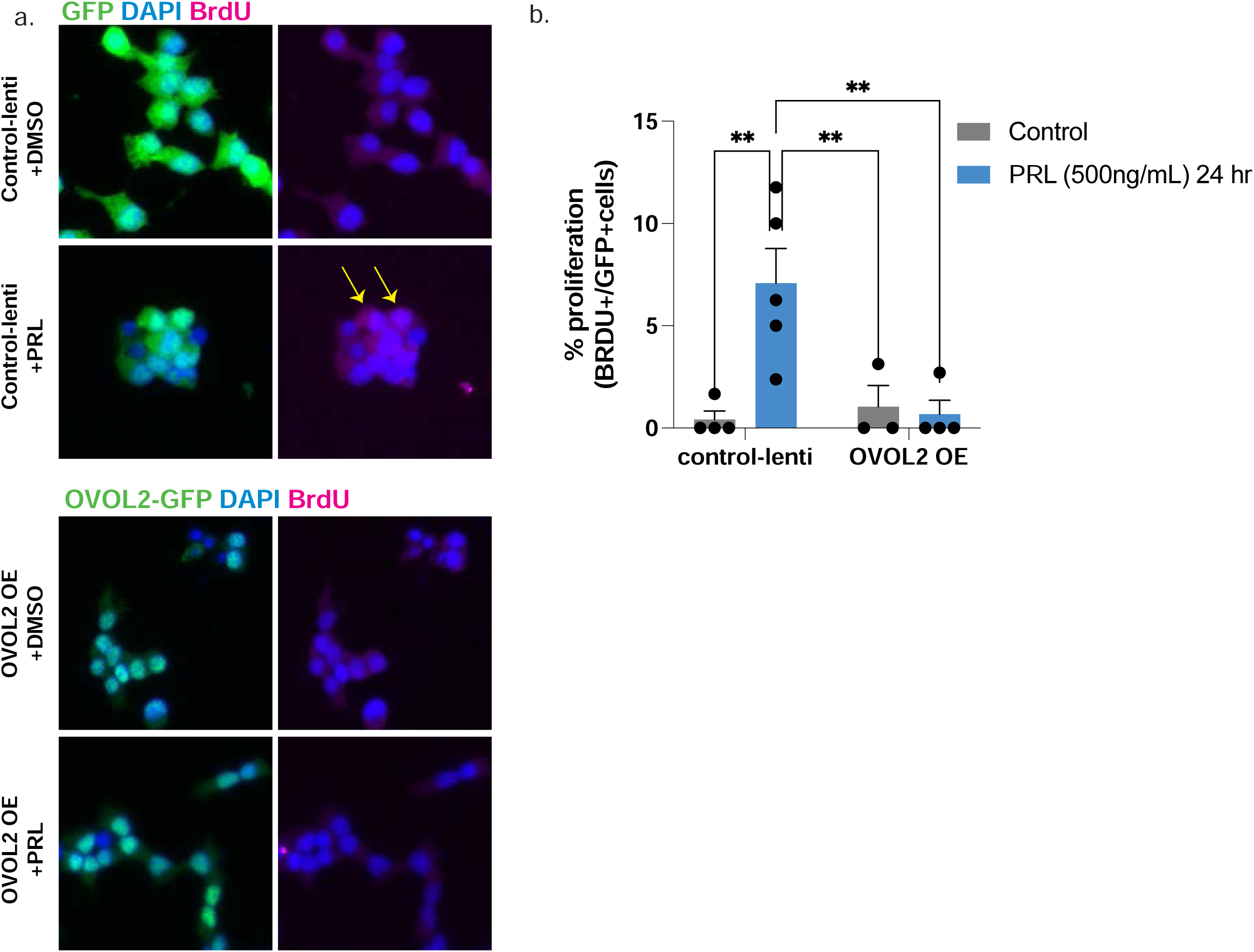
OVOL2 overexpression diminishes PRL-mediated proliferation of MIN6 cells. c. BRDU (red) detection in lentiviral infected MIN6 cells after PRL treatment. d. Quantification of BrdU incorporation of MIN6 cells after PRL treatment. Means ± SEM for N= 3 plates per condition and treatment combination. 2Way ANOVA: ** p-values <0.01

Finally, we wanted to examine whether OVOL2 expression is regulated by PRLR signaling *in vivo.* Using tissue sections from non-pregnant, pregnant and postpartum WT female mice, we observed an up-regulation of OVOL2 within β-cell nuclei of pregnant mice **(Figure 6a and 6b),** which did not persist postpartum. Confirming the requirement for PRLR signaling to induce the gestational increase in OVOL2, β-cells from pregnant β-PrlrKO did not display similar up-regulation. Taken together, our findings indicate that OVOL2 acts downstream of PRLR signaling within β-cells where it acts as a negative regulator of this pathway.

**Figure 6:**
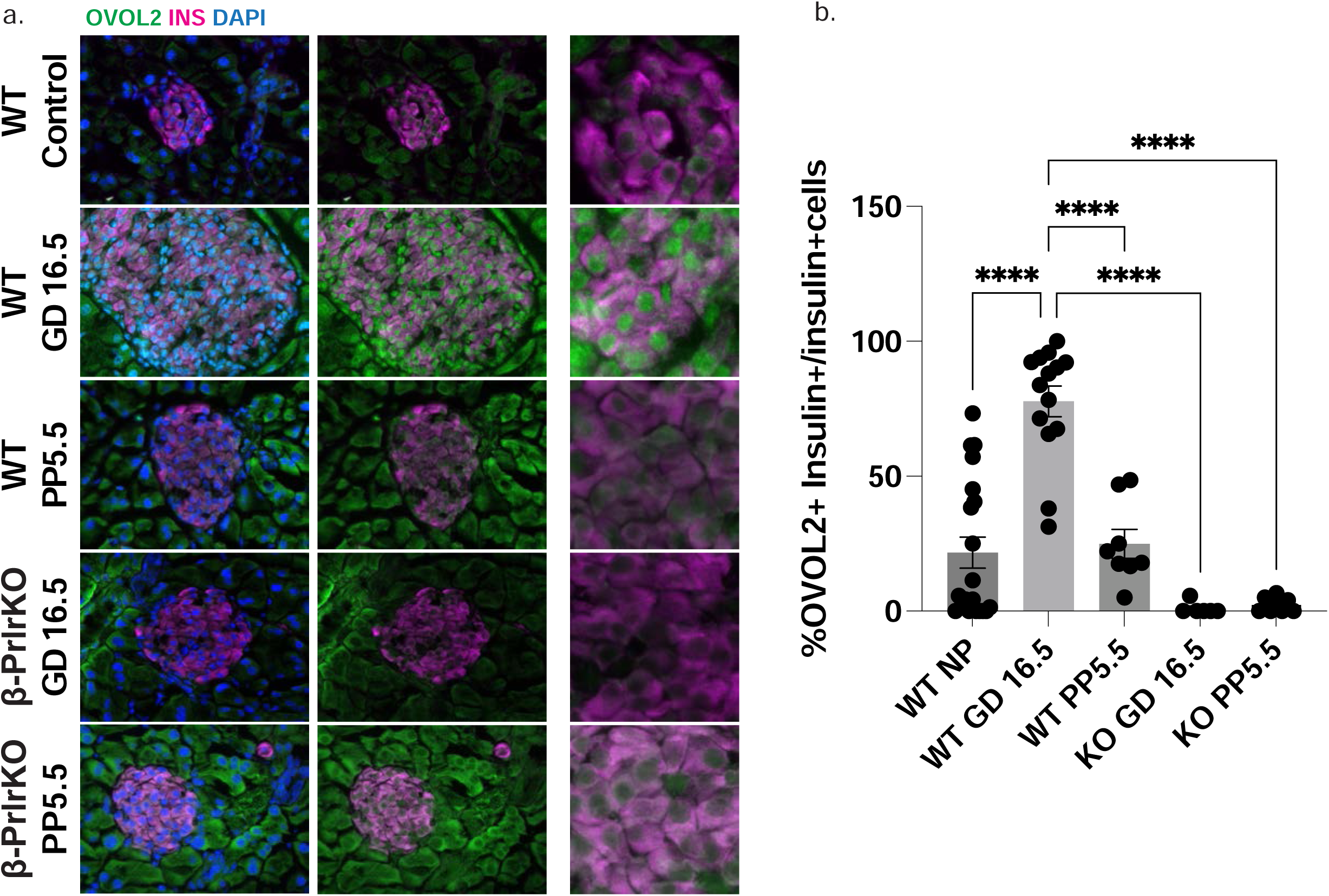
OVOL2 is found in β-cell nuclei during pregnancy but not in virgin or postpartum WT controls or in β-PrlrKO mice during pregnancy. a. Immunostaining of islets from non-pregnant, pregnancy (PREG; gestational day 16.5) or postpartum (POST; postpartum day 5) mice in control or β-PrlrKO female mice (PRLR^f/f^; RIP-Cre). OVOL2 (green), insulin (magenta) and DAPI (blue). b. Quantification of OVOL2+ and INSULIN+ cells normalized by total INSULIN+ cells. Means ± SEM for N= 2 animals per group, 10 or more islets per animal. One-way ANOVA: **** p-values <0.0001

## Discussion

Our studies demonstrate that PRLR signaling regulates changes in the chromatin accessibility of murine female β-cells. Using scATAC-seq we find PRL treatment increased chromatin accessibility within PRLR-expressing islet β-cells distinct from other islet cells which do not express high levels of PRLR. Our analysis was strengthened by including islets from β-PrlrKO animals as they show cell autonomous regulation of β-cell chromatin accessibility by PRLR.

Increases in chromatin accessibility are thought to facilitate TFs access to their binding sites, therefore facilitating changes in the cell’s transcriptome by allowing the expression of previously silenced genes. We propose that chromatin structure modifications downstream of PRLR signaling facilitates β-cell adaptations during gestation. Future studies are needed to determine how chromatin accessibility changes during gestation and postpartum both in murine and human models, to confirm this mechanism of action *in vivo*.

To determine TFs that might bind newly accessible regions after lactogen stimulation, we identified motifs enriched in DARs after PRL-treatment. Among these, we found several TFs previously reported to act downstream of PRLR signaling[7]. However, we also identified a list of novel TFs showing PRL-dependent transcriptional expression and whose motifs were overrepresented in DARs, including OVOL2. Further research is required to elucidate the temporal and dynamic regulation of these TFs, whether they mediate chromatin or other transcriptional changes downstream of PRLR, and how these contribute to islet adaptations downstream of lactogen signaling in gestation. Additionally, how OVOL2 acts downstream of PRLR to directly regulate chromatin structure in β-cells requires further investigation.

OVOL2 transcription and binding is increased upon lactogen stimulation in islets. Interestingly, although the overexpression of OVOL2 showed a trend towards enhanced PRL-dependent expression of its target gene, *Snai1,* as expected, OVOL2 overexpression also inhibited PRL-induction of *Myc* and *Rhoj,* both of which are regulators of cell proliferation[21]. Thus, overall, OVOL2 overexpression impairs PRL-dependent proliferation suggesting an inhibitory effect of OVOL2 in β-cells.

Previously, OVOL2 has been shown to inhibit tumor metastasis and act via *Myc* to promote transient proliferation in keratinocytes[21]. Similarly, OVOL2 regulates mammary morphogenesis and regeneration[44]. Although PRL treatment in islets *ex vivo* requires several days to induce β-cell proliferation[45, 46], we propose that the PRL-dependent chromatin changes we observed after only 24 hours of prolactin stimulation allow pioneer TFs to access chromatin sites to permit further adaptations during the gestational period. Chromatin accessibility studies early after lactogen treatment might identify pioneer TFs responsible for chromatin opening. We propose that OVOL2 acts as a molecular negative feedback loop downstream of PRLR signaling to prevent uncontrolled proliferation. Our immunostaining studies demonstrate that OVOL2 is expressed at GD16.5, a late pregnancy stage where β-cell proliferation is elevated but beginning to subside and might represent part of this molecular “brake” machinery as the islet prepares for the rapid reversal of proliferation in late gestation towards regression of β-cell mass in the early postpartum period. Future mechanistic interrogation of OVOL2 in pregnancy and postpartum will be critical in confirming it OVOL2 regulates β-cells *in vivo* consistent with its activity in our *ex vivo* studies of islets.

## Methods

### Animals

All procedures relating to animal care and treatment conformed to Johns Hopkins University Animal Care and Use Committee (ACUC) and NIH guidelines where these studies were performed. Animals were under direct supervision of the staff veterinarian and were monitored daily by Animal Care Staff. In addition to the sterile environment and weekly cage changes, mice were kept in a temperature and humidity-controlled room with 12-hour light/dark cycle. All mice had free access to food and water. All experiments were performed using 8– to 14-week-old virgin female mice. Wild type C57BL6J mice were obtained from Jackson Laboratory (Strain #000664). *Rip-Cre* mice[47] were crossed with *Prlr^f/f^* to create β-cell specific deletion of PRLR (β-PrlrKO), as previously reported[19].

### Islet isolations

Islets were perfused with Collagenase Type XI (Millipore Sigma, # C7657) dissolved in Hank’s Buffered Saline Solution (HBSS) (Thermo Fisher Scientific, #14175095) at 0.8 mg/mL into the common bile duct. Tissues were incubated at 37°C, washed twice in 10%FBS (Thermo Fisher Scientific, # 26140079) in HBSS and once in HBSS and subjected to discontinuous gradient using histopaque (6:5 Histopaque 1119: Histopaque 1077; Sigma-Aldrich, #11191 and #10771) and HBSS. Islet layers were handpicked in 2% FBS in HBSS for downstream assays.

### Islets culture and prolactin treatment

Isolated islets were cultured in RPMI-1640 (Thermo Fisher Scientific, # 22400089) supplemented with 10% FBS and 2% penicillin-streptomycin overnight at 37°C in a humid chamber to allow for recovery. Islets were treated with 500ng/mL of recombinant mouse prolactin (Bio-Techne, # 1445-PL-050) for 24 hours.

### scATAC-sequencing

Single cell suspensions were obtained after incubating islets in 0.05% trypsin-EDTA (Invitrogen, #25300054) at 37°C with gentle trituration using a P1000 pipette for 10 minutes. Digestion was interrupted using FBS and cells were immediately filtered using a 40um cell strainer. After a PBS wash, cells were resuspended in 0.04% BSA in PBS 1X. Cell viability and concentration were quantified using Cell Countess II (Thermo Fisher Scientific). After nuclei extraction, transposition, GEM generation and library preparation was prepared according to the manufacturer’s instructions using Chromium Next GEM Single Cell ATAC Kit (10X Genomics, #1000406), Chromium Next GEM Chip H Single Cell Kit (10X Genomics, #1000162), Single Index Kit N Set A, (10X Genomics, #1000212) and the Chromium Controller (10X Genomics Model GCG-SR-1). The Single Cell & Transcriptomics Core (SCTC, Johns Hopkins University, School of Medicine, Baltimore, MD) assessed the quality of the constructed library, performed Illumina sequencing using NovaSeq X and aid in the pre-processing of the cells using the 10X Genomics Cell Ranger pipeline.

### RNA-sequencing

RNA was isolated using the TRIzol (Thermo Fisher Scientific, #15596026). A total RNA of 500 ng was used for library construction with the NEBNext Ultra II RNA Library Prep Kit for Illumina (BioLabs, #E7775), together with the NEBNext Poly(A) mRNA Magnetic Isolation Module (BioLabs, #E7490) and the NEBNext Multiplex Oligos for Illumina (Dual Index Primer Set I, Biolabs # E7600). All preparations were performed according to the manufacturer’s instructions. The quality of the constructed library was assessed, and sequencing was performed in the Illumina Novaseq X Plus Lane system by the Genetic Resources Core Facility (GRCF, Johns Hopkins University, School of Medicine, Baltimore, MD).

### Bioinformatic analysis

#### RNA-sequencing

We performed the genome alignment using histat2 v2.2 and the output sam file was converted, sorted and indexed using samtools v1.21. A matrix of the raw counts was created using featureCounts (subread-2.1.0). To determine PRL-DEGS we used the DESeq2 package for R v. 1.46.0.

#### scATAC-sequencing

We used the Seurat and Signac packages (See table 2) to perform cell clustering, quantitative control exclusions and all the analyses presented in Figure 2 and associated supplemental tables as previously described[48–50]. We created a common set of peaks to compare all cells and confirmed transcription start site (TSS) enrichment of Tn5 insertion and followed integration of replicates and merging protocols. We calculated gene activities, using 2k base pairs(bps) upstream of the TSS, as these regions usually contain regulatory sequences, and the full gene length up a total size of 500kbps[49]. Individual cell clusters were categorized into cell types based on the gene activities of well-characterized marker genes of islet cells. We did not detect exon 5 of the *Prlr* gene in β-cells from β-PrlrKO, as opposed to β-cells from WT, consistent with efficient recombination of the floxed allele of this mouse line. To determine the motif enrichment, we used JASPAR2020[42].

### qPCR

TRIzol-based RNA extraction from whole islets was performed as previously described[8]. We used Superscript IV (ThermoFisher Scientific #18090010) for cDNA synthesis as per manufacturer’s instructions. Real-time PCR was performed with PrimeTime qPCR Probe Assays (Integrated DNA Technologies) and FastStart Universal Probe Master (Millipore Sigma #04914058001) or the primers listed in Table 1 and PowerUp SYBR Green Master Mix (Applied Biosystems #A25742) using QuantStudio™ 5 Real-Time PCR Instrument (Applied Biosystems).

**Table 1:**
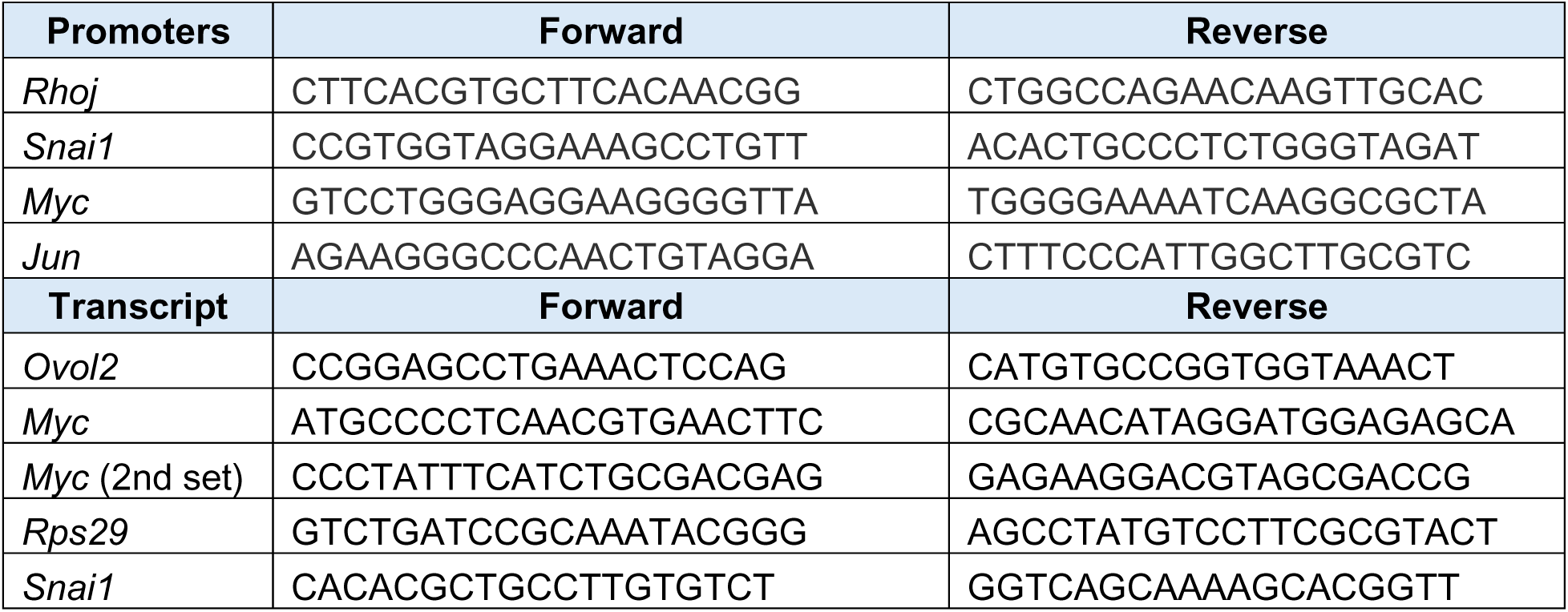
qPCR primers.

**Table 2:** Reagent or resource.

| Reagent or resource | Company-Lab | RRID or catalog # |
| --- | --- | --- |
| Rabbit anti OVOL2 (polyclonal) | Bio-Techne (Nuvous) | AB_3307612 |
| Guinea Pig anti insulin (polyclonal) | Agilent (Dako) | AB_2800361 |
| Seurat (R package) | Satija Lab | SCR_016341 |
| Signac (R package) | Stuart Lab | SCR_021158 |
| SeuratWrappers (R package) | Satija Lab | SCR_022555 |
| JASPAR2020 (R package) | [42] | RRID:SCR_003030 |
| rGreat | Zuguang Gu | RRID:SCR_028008 |
| TFBSTools | G Tan | SCR_024260 |
| pLV[Exp]-EGFP/Puro-EF1A>mCherry (Ctrl-lenti) | VectorBuilder | VB010000-9492agg |
| pLV[Exp]-Puro-CMV>mOvol2[NM_026924.3]/3xGGGGS/EGFP (OVOL2OE) | VectorBuilder | VB250505-1105ZYC |

### ChIP-qPCR

For ChIP we pooled islets from 10 mice and separated two groups to treat with PRL or DMSO after an overnight recovery. After 24 hours of treatment, islets were fixed in 1% formaldehyde, followed by 1M glycine treatment. After cell lysis, DNA was sheared using the Covaris E220 system (SCTC, Johns Hopkins University, School of Medicine, Baltimore, MD) and immunoprecipitation and DNA purification was performed using the MAGnify Chromatin Immunoprecipitation System (Applied Biosystems #492024) according to the manufacturer’s instructions. We designed custom primers targeting promoter regions of a subset of genes containing the OVOL2 binding motif and that were transcriptionally affected by PRL treatment to assess OVOL2 binding. qPCR was performed as described above.

### MIN6 cells

MIN6 cells, a mouse pancreatic β-cell line derived from insulinomas in transgenic mice(<u>CVCL_0431</u>) [43] were provided by Maria Golson (Johns Hopkins School of Medicine, Baltimore, MD). MIN6 cells were cultured in Dulbecco’s Modified Eagle’s Medium (DMEM #30–2002, ATCC, Manassas, VA) containing 15% FBS, (Thermo Fisher Scientific #26140079), 10mM HEPES (Thermo Fisher Scientific #15630080), 100units penicillin-streptomycin (Thermo Fisher Scientific #15140122) and 70μM β-Mercaptoethanol (Thermo Fisher Scientific # 21985023). Cells were kept in a humidified incubator at 37°C with 5% CO_2_.

### Lentiviral infection

MIN6 cells were infected with the corresponding lentiviral particles when they reached approximately ∼50% confluency at 1 MOI in DMEM media containing 10mM HEPES, 100units penicillin-streptomycin and 2μg/mL of polybrene (VectorBuilder). After 24 hours, the lentiviral-containing media was removed and cells were allowed to recover for 48 hours in regular DMEM media, as described above. Selection was performed using puromycin treatment daily for 3-days, and cells were allowed to recover for an additional 2 days before additional experiments were performed.

### Immunofluorescence

Mouse pancreata were fixed for 30 min at room temperature in 4 % paraformaldehyde (PFA, Cat # 100503-917, VWR, Radnor, PA). Fixed tissues were cryoprotected using a sucrose solution and then embedded in OCT compound (Cat # 23-730-571, Fisher Scientific, Hampton, NH), for sectioning. Slides were washed in PBS, permeabilized in 0.2% TritonX-100 and blocked for one-hour at room temperature in 3 %BSA (Millipore Sigma # A9647) and 5 % normal goat serum (Cell Signaling Technologies #5425) in a humidified chamber for 1-hour at room temperature. Primary antibodies diluted in blocking solution were incubated overnight at 4°C in a humidified chamber. Secondary antibodies and DAPI were incubated for 1-hour at room temperature. After washes, sections were mounted using Fluoromount Aqueous Mounting Medium (Sigma Aldrich #F4680). Images were captured using a Lionheart FX automated cell imaging microscope (Agilent-BioTek, Santa Clara, CA). Fluorescent quantification was performed using ImageJ software (V 2.16.01.54p).

### Quantification and statistical analysis

Statistical and graphing analyses were performed using GraphPad Prism software (V 10.5.0) or using R (V 4.4.2) through RStudio (C 2025.05.0). For one-way ANOVA, post hoc testing was performed with Tukey’s multiple comparisons test. Statistical testing performed and statistical significance are specified within figure legends. Statistical analyses were based on at least 3 independent experiments and described in figure legends.

## Data availability

All data needed to evaluate the conclusions in the paper are present in the paper and/or the Supplementary Materials. The scATAC-seq and RNA-seq data generated during the current study are available in the NCBI GEO repository under the accessions GSE319576 and GSE319525 respectively.

