## Supplemental figures for "Single-cell ATAC-seq Reveals OVOL2 as a Downstream Negative Regulator of PRL-Mediated Chromatin Accessibility"

### Supplemental Figure legend

**Supplemental figure 1: scATAC-sequencing of islets show that PRLR signaling mediates DARs in β -cells**

a. Uniform Manifold Approximation and Projection (UMAP) of scATAC-seq profiles from islet cells. Individual dots represent single cells, color-coded by Genotype, Treatment or Sample.
b. qPCR of islets for *Tph1* and *Tph2* after PRL-treatment. Relative mRNA levels were normalized to *Rps29* and calculated using the $2^{-DDCT}$ method. Student t-test: ** p value <0.01.
c. Heatmap of cluster-defining cell marker peaks. Normalized accessibility signal (expression) for known islet cell markers per cluster. Data are grouped by cell type; color intensity represents relative accessibility.
d. Distance-to-TSS profiles for scATAC-seq peaks within β-cells. The x-axis represents the genomic distance (kbp) from the peak center to the TSS, and the y-axis shows the relative fraction of total peaks.

**Supplemental figure 2: Visualization of chromatin accessibility within PRL-DEGs**

a. Coverage plots from β-cells show reduction of sequencing reads belonging to exon 5 of the *Prlr* gene in β-PrlrKO but not in WT regardless of PRL treatment
b. Coverage plots from β-cells showing differential chromatin accessibility in β-cells from β-PrlrKO and WT regardless of PRL treatment

**Supplemental figure 3: Identification TFs which are PRL-DEGs and with overrepresented motifs in WT β-cells PRL-DARs**

a. Enhanced volcano plot showing transcriptional differences identified by RNA-sequencing (left). Term-Gene graph shows DEGS involved in significantly enriched terms in islets treated with PRL (term-gene created using pathfindR) (right).
b. Motif plot for the STAT family of transcription factors overrepresented after PRL-treatment in WT β-cells

**Supplemental figure 4: PRL-dependent binding and gene regulation by OVOL2**

a. ChIP-qPCR for the *Jun* promoter from islets treated with PRL. Binding to promoters was calculated in relation to input (% input) and evaluated against rabbit IgG as a control for non-specific binding. Means ± SEM for N= 5 experiments: 2Way ANOVA: ** p value <0.01.
b. qPCR evaluation of MIN6 cells infected with control (Ctrl-lenti) or OVOL2OE after PRL treatment. Relative mRNA levels were normalized to *Rps29* and calculated using the $2^{-\Delta\Delta CT}$ method. Means ± SEM for N= 3 plates per condition and treatment combination. 2Way ANOVA: * p value <0.05 and ** p value <0.01.

#### Supplemental Fig. 1

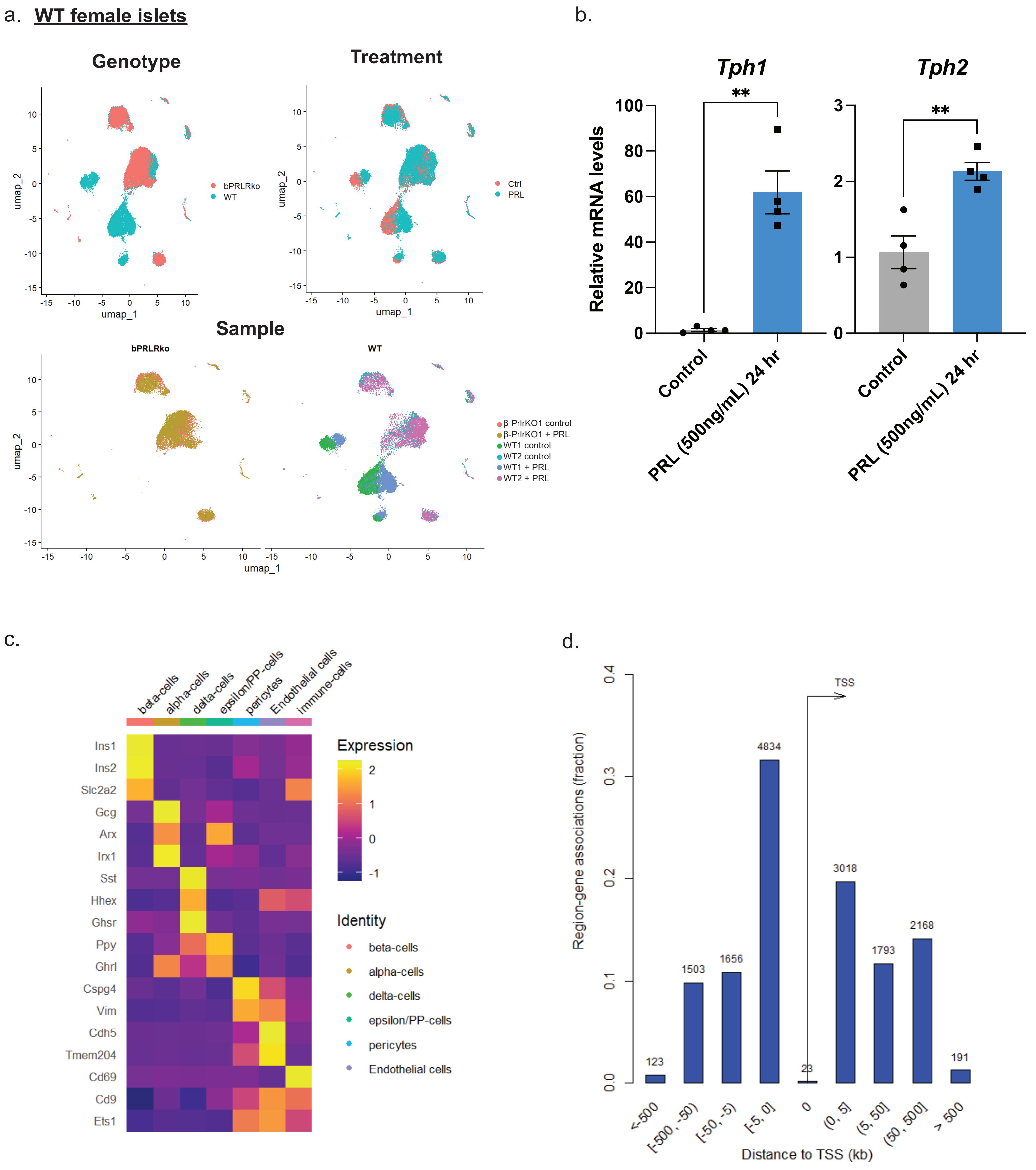

**Supplemental Fig. 2**

a. β-cells

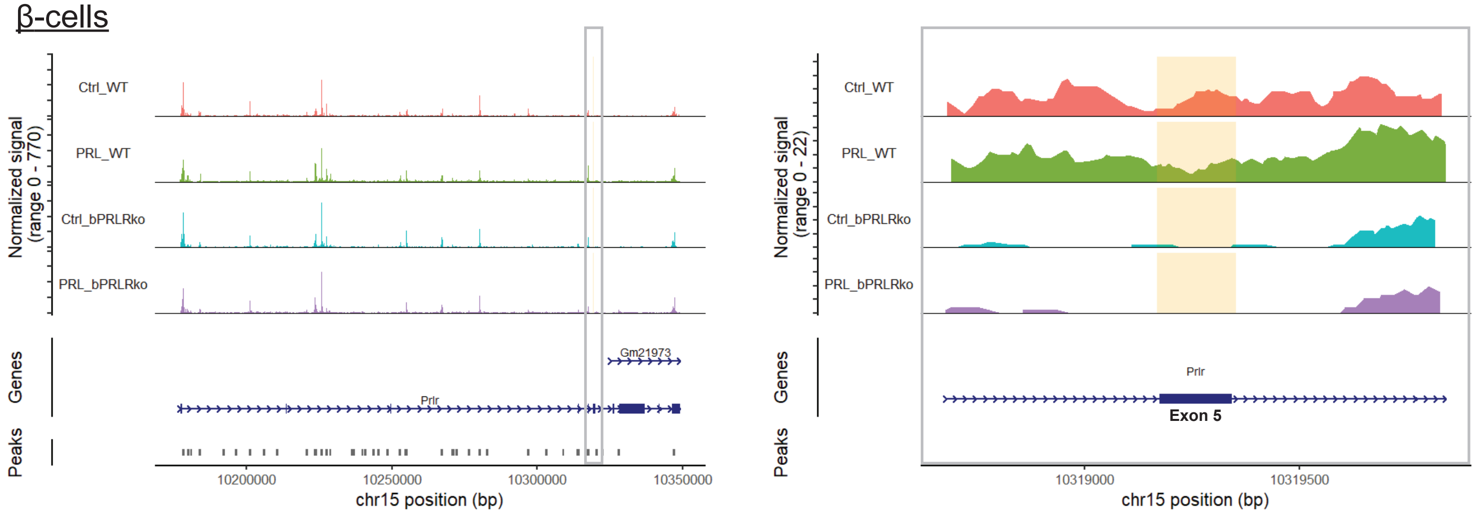


b. β-cells

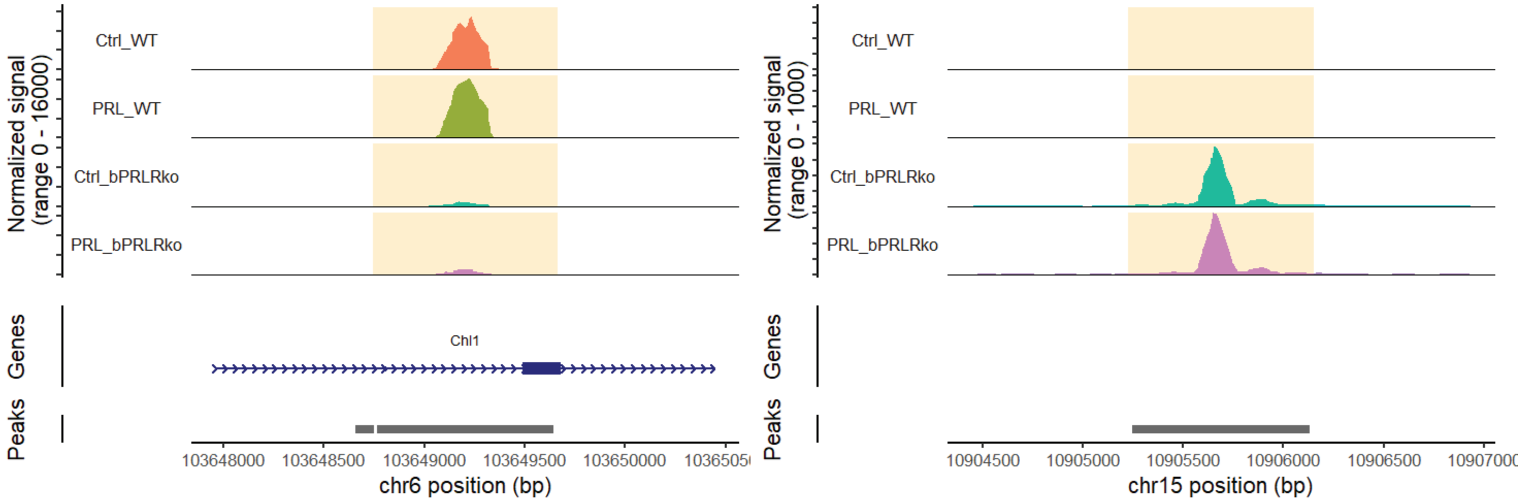

**Supplemental Fig. 3**

a.

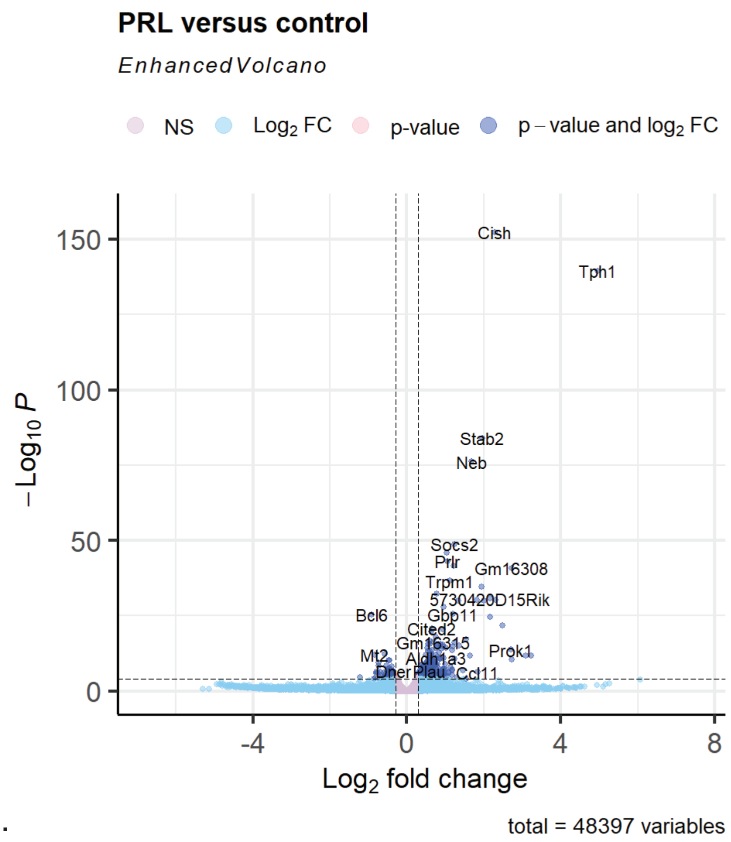


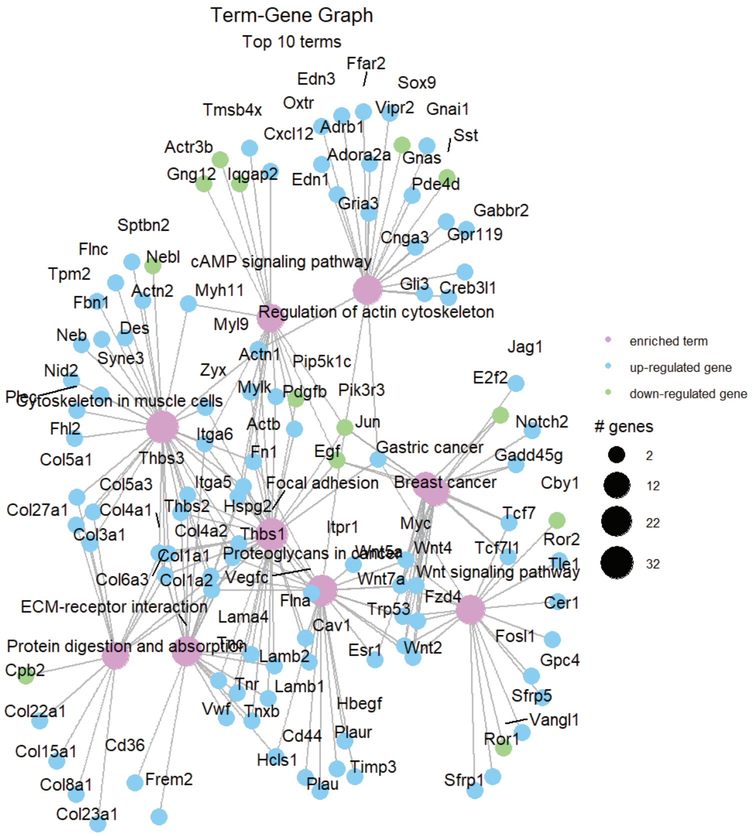


b.

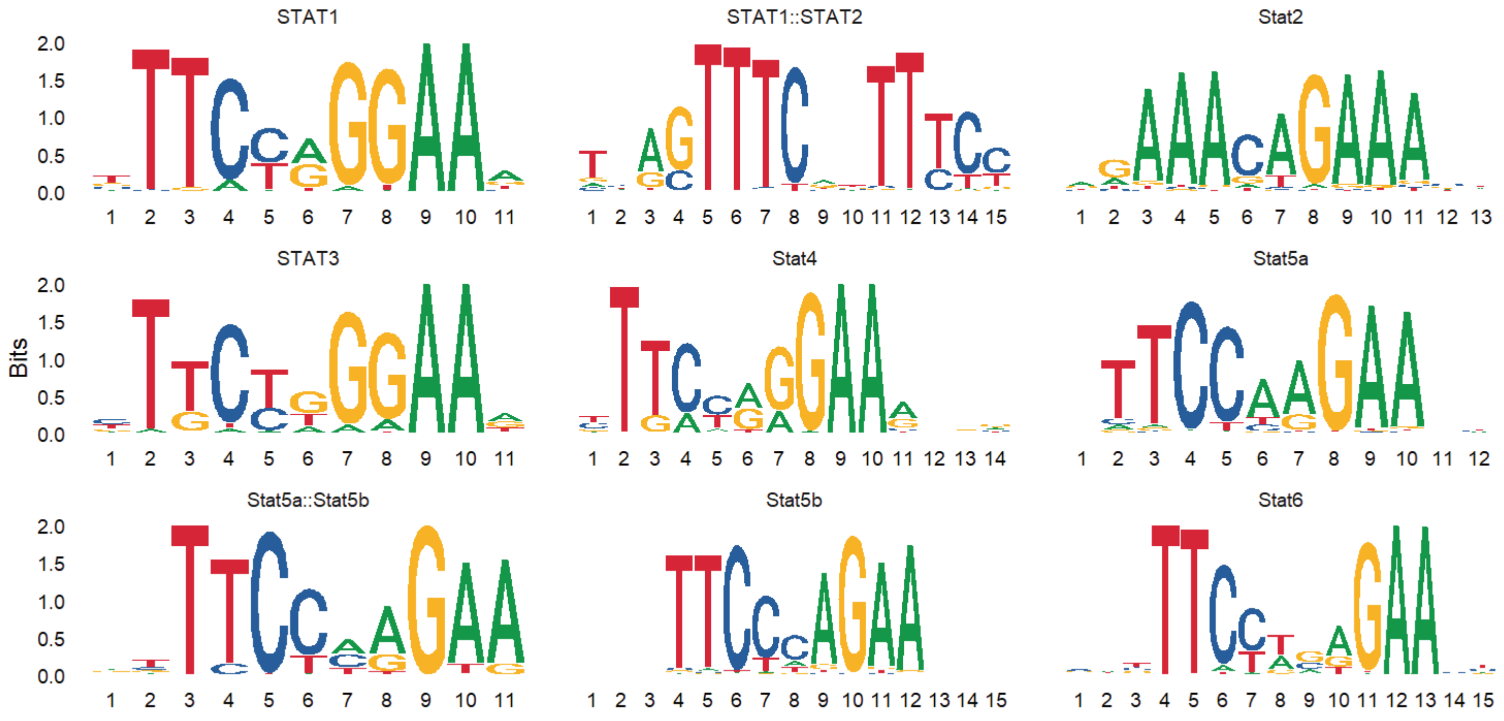

Supplemental Fig. 4

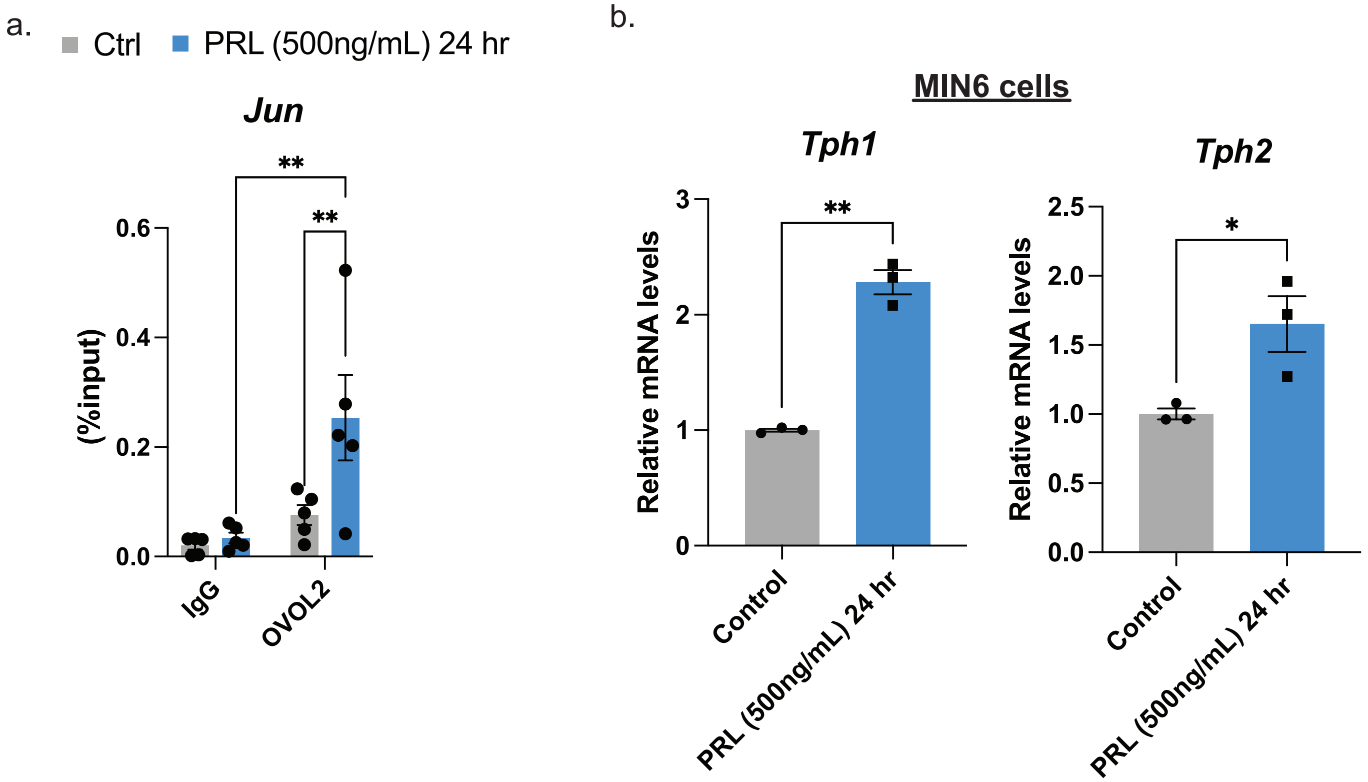
